# Glucose Transporter Expression and Regulation Following a Fast in the Ruby-throated Hummingbird, *Archilochus colubris*

**DOI:** 10.1101/2020.06.12.148619

**Authors:** Raafay S. Ali, Morag F. Dick, Saad Muhammad, Dylan Sarver, G. William Wong, Kenneth C. Welch

## Abstract

Hummingbirds subsist almost exclusively on nectar sugar and face extreme challenges blood sugar regulation. Transmembrane sugar transport is mediated by facilitative glucose transporters (GLUTs) and the capacity for sugar transport is dependent on both the activity of GLUTs and their localisation to the plasma membrane (PM). In this study, we determined the relative protein abundance in whole-tissue (WT) homogenates and PM fractions via immunoblot using custom antibodies for GLUT1, GLUT2, GLUT3, and GLUT5 in flight muscle, heart, and, liver of ruby-throated hummingbirds (*Archilochus colubris*). GLUTs examined were detected in nearly all tissues tested. Hepatic GLUT1 was minimally present in WT homogenates and absent in PM fractions. GLUT5 was expressed in hummingbird flight muscles at levels comparable to that of their liver, consistent with the hypothesised uniquely high fructose-uptake and oxidation capacity of this tissue. To assess GLUT regulation, we fed ruby-throated hummingbirds 1M sucrose *ad libitum* for 24 hours followed by either 1 hour of fasting or continued *ad libitum* feeding until sampling. We measured relative GLUT abundance and concentrations of circulating sugars. Blood fructose concentration in fasted hummingbirds declined from ∼5mM to ∼0.18mM, while fructose-transporting PM GLUT2 and PM GLUT5 did not change in abundance. Blood glucose concentrations remained elevated in both fed and fasted hummingbirds, at ∼30mM, while glucose-transporting PM GLUT1 and PM GLUT3 in the flight muscle and liver, respectively, declined in fasted birds. Our results suggest that glucose uptake capacity is dynamically reduced in response to fasting, allowing for maintenance of elevated blood glucose levels, while fructose uptake capacity remains constitutively elevated promoting depletion of blood total fructose within the first hour of a fast.

**Summary statement:** Hummingbird ingest nectar rich in glucose and fructose. When fasted, tissue capacity for circulating glucose import declines while remaining elevated for fructose. This may underlie maintenance of high blood glucose and rapid depletion of blood fructose.

## Introduction

Hummingbirds primarily subsist on a diet of floral nectar high in sucrose, glucose, and fructose (del Rio et al., 1992). They are capable of oxidising glucose, fructose, or both, to power their characteristic hovering behaviour (Chen and Welch, 2014). When blood sugar concentrations are elevated, hummingbirds rely exclusively on these exogenous sugars to fuel nearly all the metabolic needs of their active cells (Welch et al., 2018). As such, they exhibit remarkable adaptations that enhance both the capacity for immediate rapid uptake and metabolism and the long-term storage of these sugars (Price et al., 2015; Welch et al., 2018). When possible, circulating sugars are incorporated into hummingbirds’ fat stores through *de-novo* lipogenesis by their liver (Suarez et al., 1988). As hummingbirds enter periods of hypoglycaemia, such as sleeping or fasted states, the entirety of their metabolic fuel source switches from circulating sugars to triglycerides derived from these fatty-acid stores (Eberts et al., 2019; Suarez et al., 1990). This switch is rapid, and a transition back to sugar metabolism occurs within a few minutes of sugar ingestion (Suarez and Welch, 2017). Furthermore, the switch from reliance on lipid oxidation to carbohydrate oxidation is nearly complete, such that mixed fuel-use does not occur for very long in hummingbirds with access to sufficient floral nectar (Welch et al., 2018).

Hummingbird digestive physiology facilitates rapid sugar transport across the intestinal lumen and into circulation (Karasov, 2017). A high cardiac output and capillary-to-muscle-fibre ratio ensures high transport capacity of sugars to the site of active cells (Mathieu-Costello et al., 1992; Suarez, 1992). Sugars are then facilitatively imported across the plasma membrane (PM) of active cells (Suarez and Welch, 2011). Here, *in-vitro* studies of hummingbird muscle cells have demonstrated that the phosphorylation capacity of cytosolic kinases for glucose appears sufficient in providing energy for sustained hovering, although this may not be true for fructose (Myrka and Welch, 2018). As both delivery to and phosphorylation of glucose within muscles operate at rates near the theoretical maximum in vertebrates (Suarez et al., 1988; Suarez and Welch, 2017) it is likely that regulation at the site of import itself exerts a great deal of control over the flux through the entirety of the sugar oxidation cascade. Along with delivery and phosphorylation, the sugar import step is a rate-limiting process in the paradigm outlined by Wasserman et al. (2011) and is nearly entirely dependent on the presence and distribution of active glucose transporters (GLUTs) (Wasserman, 2009). These proteins are a family of transmembrane solute transporters (Mueckler and Thorens, 2013).

Studies of mammalian GLUTs demonstrate that their expression in the PM is regulated by a variety of intra- and extracellular factors, including blood sugar and insulin concentrations, exercise, and stress (Egert et al., 1999; Guma et al., 1995; Yang and Holman, 1993). The expression and functional distribution and regulation of hummingbird GLUTs, however, remains relatively unknown. Studies on GLUT isoforms of the closest relatively well-examined avian species, the chicken (*Gallus gallus domesticus*), are fragmented and the distribution of avian GLUT isoforms is not fully understood (Byers et al., 2018; Suarez and Welch, 2011; Sweazea and Braun, 2006). It is known that chicken GLUT1 and GLUT3 share sequence homologies of ∼80% and ∼70%, respectively, with human GLUTs, but other isoforms such as GLUT2 and GLUT5 only share ∼65% and ∼64% sequence homology (calculated via NCBI BLAST (Boratyn et al., 2012), summarised in Table S6). It is also clear that they are regulated very differently in each class (Wagstaff and White, 1995; Yamada et al., 1983). Despite this, the literature on mammalian GLUTs provides a useful foundation for understanding the affinities and ligand-specificity of avian, including hummingbird, GLUTs. In mammals, GLUT3, followed by GLUT1, show the highest affinities for glucose; *K*_*m*_ ≈ 1.5mM (Thorens and Mueckler, 2010) and *K*_*m*_ ≈ 3-5mM (Zhao and Keating, 2007), respectively. GLUT5 transports fructose (*K*_*m*_ ≈ 11-12mM; Douard and Ferraris, 2008), and is largely found in mammalian enteric and renal tissue (Douard and Ferraris, 2008), although some presence in hepatic tissue has also been noted (Godoy et al., 2006; Zhao et al., 1993). GLUT2, uniquely, shows affinity for both sugars. While its affinity for glucose and fructose (*K*_*m*_ ≈ 17mM and *K*_*m*_ ≈ 76mM, respectively; Zhao and Keating, 2007) is relatively low compared to other isoforms, it plays a dominant role in hepatic sugar transport (Wood and Trayhurn, 2003).

Importantly, it is only when GLUT isoforms are expressed and active in the PM that transmembrane sugar transport can occur from the blood into the active cell (Guma et al., 1995; Wasserman, 2009; Yamada et al., 1983). In mammals, GLUT4 translocation to the PM by insulin-stimulation following feeding is known to recruit other GLUT isoforms to the PM as well, increasing the sugar import rate into active cells (Guma et al., 1995). Hummingbirds (Welch et al., 2013), much like chickens (Byers et al., 2018), do not express transcript or protein of the insulin-sensitive GLUT4 isoform. Chicken insulin levels do not significantly change with dietary status (Simon et al., 2011), and this is presumably also true in hummingbirds. Further, circulating insulin does not significantly increase sugar import in chicken muscles (Chen, 1945), though it may in the liver (Dupont, 2009; Zhang et al., 2013). Lastly, and unlike mammals, hummingbirds have limited intramuscular glycogen stores (Suarez et al., 1990), and therefore rely on newly imported sugars from circulation for carbohydrate oxidation (Welch et al., 2018). Despite missing critical elements of the insulin-GLUT4 pathway, fed hummingbirds utilise circulating sugars, when available, at very high rates to meet their metabolic demands (Suarez and Welch, 2017).

Previous studies have confirmed the presence of GLUT1 and GLUT5 transcript in nearly all hummingbird tissue examined (Myrka and Welch, 2018). Immunohistochemistry of hummingbird myocytes using a commercial antibody for GLUT1 have also shown GLUT1 localisation to the PM (Welch et al., 2013), though, the results were not definitive. In this study, using custom antibodies for the different isoforms of hummingbird GLUTs, we sought to identify the tissue-specific protein distribution and to quantify the abundance in the PM, of GLUT1, GLUT2, GLUT3, and GLUT5. We predicted GLUT1 would be detected in hummingbird flight muscle, cardiac, and liver tissue, in accordance with its ubiquitous presence in mammalian tissue (Mueckler and Thorens, 2013), as well as its previous detection in hummingbird myocytes (Welch et al., 2013). As GLUT2 plays a stronger role in enteric (Karasov, 2017) and hepatic (Mueckler and Thorens, 2013) sugar transport, we predicted that its abundance would be limited in muscles and more predominantly found in the liver. In mammals, GLUT3 is observed in close association with GLUT1 (Simpson et al., 2008) and may function as a replacement for GLUT4 in certain muscle developmental stages (Klip et al., 1996). We expected to detect GLUT3 in tissues also expressing GLUT1. We also expected to find GLUT5 in both the liver and muscles, as hummingbird muscles are capable of supporting hovering flight on fructose-only meals (Chen and Welch, 2014). To further characterise the regulatory aspects of hummingbird GLUTs, we compared the abundance of GLUT1, GLUT2, GLUT3, and GLUT5 in the PM of fed and fasted hummingbirds. We also measured levels of circulating glucose and fructose in these birds. Based on previous measurements of hummingbird blood glucose (Beuchat and Chong, 1998), we expected to see high levels of glucose (∼40mM) in the fed condition and lower levels in the fasted (∼15mM). Previous measurements of hummingbird blood fructose have not been made. However, similar to that of frugivorous bats (Keegan, 1977), we predicted blood fructose concentrations in fed hummingbirds to be ∼5-10mM in fed and ∼0mM in fasted hummingbirds. Given the rapid switching between glucose or fructose oxidation and oxidation of lipid stores in foraging versus fasting hummingbirds, we expected a greater abundance of PM GLUT1, PM GLUT3, and PM GLUT5 in flight muscle and liver of fasted hummingbirds. Finally, we expected little difference in between GLUT2 abundance in the PM of tissue from fed and fasted hummingbirds.

## Materials and Methods

### 1.1 Animal Use and Ethics Statement

This study was approved and performed adhering to the requirements of the University of Toronto Laboratory Animal Care Committee and the Canadian Council on Animal Care. Twelve adult male ruby-throated hummingbirds (*Archilochus colubris*) were captured in the early summer at the University of Toronto Scarborough (UTSC) using modified box traps and housed individually in Eurocages (Corners Ltd, Kalamazoo, MI, USA) in the UTSC vivarium. They were provided with perches and fed on a maintenance diet of NEKTON-Nectar-Plus (Keltern, Germany).

All hummingbirds were provided with a sucrose solution for 24 hours prior to the experiment. Birds were divided into a fed group (n = 6), which was provided with *ad-libitum* 1M sucrose solution up to sampling, beginning at 10AM, and a fasted group (n = 6), which was deprived of any food for a 1 hour duration prior to the 10AM sample collection. To minimize interindividual variation in activity level and energy expenditure, birds from both treatment groups were held in small glass jars, perched on wooden dowels, in which the were constrained from flying, for the duration of the 1 hour fast. Respirometry measurements by Chen and Welch (2014) have previously shown that this is sufficient time for the fasted hummingbirds to shift from fuelling metabolism with circulating sugars to fats. Fed hummingbirds will continue to exclusively metabolise sugars. Hummingbirds were then anaesthetised with isofluorane inhalation and euthanized via decapitation. Immediately after decapitation, blood was sampled from the carotid artery using heparinized capillary tubes and spun at 3800 *g* for 10 minutes at room temperature and the plasma stored at -80 °C. Flight muscle (the pectoralis and supracoracoideus muscles), heart, and liver were extracted and frozen with isopentane cooled with liquid nitrogen. All tissues were stored at -80°C.

### 1.2 Circulating Sugar and Metabolite Analysis

Plasma samples were sent to the Metabolomics Innovation Centre (TMIC) at the University of Victoria (Victoria, British Columbia, Canada) to be analyzed via service 45 (absolute quantitation of central carbon metabolism metabolites and fructose) found here: https://www.metabolomicscentre.ca/service/45. Quantitation of glucose and fructose concentrations in plasma samples was achieved via chemical derivatization – liquid chromatography – multiple reaction monitoring/mass spectrometry (LC-MRM/MS) following a protocol outlined by Han et al. (2016). Quantitation of central carbon metabolites (organic acids; lactate and pyruvate) was done via the protocol outlined by Han et al. (2013).

### 1.3 Antibody Design, Production, and Isoform Specificity

Anti-rabbit polyclonal antibodies for GLUT isoforms were designed in conjunction to minimise cross-reactivity using the services of Pacific Immunology (Ramona, CA, USA). Epitope design was accomplished using messenger RNA (mRNA) sequences for ruby-throated hummingbird GLUT isoforms 1, 2, 3, and 5 that were obtained from the hummingbird liver transcriptome (Workman et al., 2018). The concentration of the affinity-purified antibody samples was assessed using ELISA by Pacific Immunology (ab-GLUT1 ≈ 1.1 mg·ml^-1^, ab-GLUT2 ≈ 5.7 mg·ml^-1^, ab-GLUT3 ≈ 2.6 mg·ml^-1^, ab-GLUT5 ≈ 1.0 mg·ml^-1^). The final experimental dilutions were determined empirically through preliminary experiments and are provided below.

#### 1.3.1 Generation of mammalian expression plasmids encoding *A. colubris* GLUT1, GLUT2, GLUT3, and GLUT5

The cDNA encoding *A. colubris* GLUT1 (NCBI Accession Number MT472837), GLUT2 (MT472838), GLUT3 (MT472839), and GLUT5 (MT472840) were synthesized by GenScript based on the full-length mRNA sequences derived from our previously published RNA sequencing data (Workman et al., 2018). The V5 epitope tag (encoding the peptide “GKPIPNPLLGLDST”) was inserted at the 3’ end of each cDNA immediately after the last coding amino acid. All epitope-tagged cDNA sequences were cloned into the EcoRI restriction site of the mammalian expression vector, pCDNA3.1 (+) (Invitrogen). All expression plasmids were verified by DNA sequencing.

#### 1.3.2 Specificity immunoblots

SDS-PAGE was run on cell lysates of HEK293T cells transiently transfected, using lipofectamine 2000 (Invitrogen), with hummingbird GLUT1, GLUT2, GLUT3, or GLUT5 (acGLUT1, GLUT2, GLUT3, or GLUT5) expression vectors; all containing a V5 tag. Cell lysates produced using RIPA buffer (50 mM Tris-HCl, pH 7.4; 150 mM NaCl; 1 mM EDTA; 1%Triton X100; 0.25% deoxycholate) supplemented with protease and a phosphatase inhibitor cocktail (MilliporeSigma, Burlington, Massachusetts, USA and Roche, Basel, Switzerland; respectively). Each lysate was confirmed to express the appropriate recombinant protein at the expected size using an anti-V5 antibody produced in rabbit (Sigma V8137). Isoform specificity was tested via immunoblotting all cell lysates (empty vector control, acGLUT1, GLUT2, GLUT3, and GLUT5) with each novel acGLUT antibody and observing GLUT protein signal overlap; none was observed. Briefly, each immunoblot lane represents a cell lysate produced from an entire well of a 6-well cell-culture dish (Thermo Scientific, Nunc). Lyates were diluted with SDS loading dye (final concentration: 50 mM Tris-HCl, pH 7.4, 2% SDS, 6% glycerol, 1% 2-ME, and 0.01% bromophenol blue) and not boiled. An equal volume of each lysate was added to the designated lane on a 12% polyacrylamide gel (Bio-Rad, Hercules, CA, USA) and separated by electrophoresis. The BioRad Trans-Blot Turbo semidry system was used to transfer protein onto PVDF membranes. Blots were blocked in 5% non-fat milk in Phosphate buffered saline with Tween 20 (PBST) and exposed to primary antibodies overnight at 4°C. After washing, blots were exposed to HRP-conjugated secondary antibody (Anti-Rabbit IgG, 7074S, Cell Signaling Technology, Danvers, MA, USA) for 1 h at room temperature and developed in ECL (Amersham ECL Select; GE Healthcare, Chicago, IL, USA). Bands were visualized with the MultiImage III FluorChem Q (Alpha Innotech, San Leandro, CA, USA). Primary antibodies were diluted 1:1000 in PBST + 0.02% sodium azide. The secondary antibody was diluted 1:10,000 in PBST + 0.02% sodium azide.

### 1.4 Tissue Sample Preparation

Each sample underwent either a plasma membrane fractionation protocol established by (Yamamoto et al., 2016) and slightly modified by replacing NP-40 (nonidet P-40) with Triton X-100 (Sigma-Aldrich, St. Louis, Missouri) to obtain only PM-proteins, or a radioimmunoprecipitation assay buffer (RIPA) homogenisation (part of the same protocol) to obtain all proteins contained in a whole-cell. Fractionation used different detergent concentrations (0.1%, 1%, 2%) in the homogenisation buffers to solubilise proteins and create protein-detergent complexes depending on whether they are in the hydrophilic (cytosolic) domain or the hydrophobic (PM) domain.

#### 1.3.3 Buffer composition

Buffer A01 (0.5M DTT, ddH_2_O, and 0.1% v/v Triton X-100), A1 (0.5M DTT, ddH_2_O, and 1% v/v Triton X-100), and 2× RIPA (20mM Tris-HCl, pH 8.0, 300mM NaCl, 2% v/v Triton X-100, 1% w/v sodium deoxycholate, 0.2% w/v sodium dodecyl sulfate (SDS), 1mM DTT) were prepared. All reagents were cooled to 4°C before homogenisation and included Sigma P8340 protease inhibitor cocktail.

#### 1.3.4 Homogenisation and plasma membrane fractionation

20 mg of flight muscle, liver, or heart was cut on a cold aluminum block and immediately placed in an ice-bath. The tissue was minced in buffer A01 with scissors and homogenised using a VWR handheld pestle homogenizer (BELAF650000000) The homogenate was passed through a 21G needle three times to liberate nuclear and intracellular proteins. An aliquot of the homogenate was left on ice for 60 minutes in 2× RIPA buffer. This whole-tissue RIPA-fraction was then centrifuged at 12,000*g* for 20 minutes at 4°C, allowing proteins to be solubilised. The supernatant was collected and stored at -80°C as the whole-tissue (WT) homogenate. The remainder of the homogenate was centrifuged at 200*g* for 1 min at 4°C. The upper phase was set aside, and 90µL of buffer AO1 was added to the lower phase which was homogenised for 10s. The lower phase was centrifuged at 200*g* for 1 minute and added to the tube containing the upper phase. The combined phases were centrifuged at 750*g* for 10 minutes. The supernatant consisting of non-PM proteins was removed. The remainder of the protein-detergent complexed pellet was resuspended with and kept on ice for 60 minutes. After centrifugation at 12000*g* for 20 minutes, and the supernatant containing only PM-associated proteins was collected as the “plasma membrane fraction”.

### 1.5 SDS-PAGE

10% resolving and 4% stacking gels were cast using a 15-well comb and the AA-Hoefer Gel Caster Apparatus (10%; 33% 30%-Acrylamide (37.1:1), 33% Separating gel buffer (1.5 M Tris Cl, 0.4% SDS), 55% ddH_2_O, 0.65% ammonium persulfate (APS), 5.5% TEMED), (4%; 13.4% 30%-Acrylamide, 9.3% Stacking gel buffer (0.5 M Tris Cl, 0.4% SDS), 33% ddH_2_O, 0.06% APS, 3.3% TEMED). Samples were incubated in a 1:1 (w/v) ratio of 2× sample buffer (0.2M DTT, BioRad Laemmli Sample Buffer #1610737) at room temperature for 20 minutes. The AA-Hoefer SE600 Vertical Gel Electrophoresis apparatus was set up with 6L running buffer (10% BioRad 10× Tris/Glycine/SDS #1610732, 90% ddH_2_O). The gel was run at 90V for 20 minutes and 110V for another 75 minutes with power supplied from an AA-Hoefer PS200HC Power Unit.

#### 1.4.1 Electroblot and immunoblot

The SDS-PAGE gel was transferred to 0.45µm pore nitrocellulose (NC) membrane (GE Life Sciences #10600003 Protran Premium 0.45 NC) using the AA-Hoefer TE22 Mighty Small Transfer unit at 110V for 90 minutes with water cooling and immersion in an icebath. The transfer buffer consisted of 192mM glycine, 24.8mM Tris, 0.00031% SDS, 20% methanol. To normalise, a total-protein stain, SYPRO Ruby Red Blot (BioRad #1703127), was used and imaged on a Bio-Rad PharosFX Molecular Imager (#1709460) using a 532nm laser and captured with a 600-630nm band pass filter. The membranes were incubated with primary antibody overnight at the following dilutions in PBST (phosphate-buffered saline, 0.1% Tween-20) buffer: GLUT1 (1:250), GLUT2 (1:2000), GLUT3 (1:2000), GLUT5 (1:500). Membranes were then incubated with anti-rabbit horseradish-peroxidase-conjugated secondary antibody (Cell Signalling Technology #7074) at 1:1000 dilution with PBST. Finally, Pierce Electrochemiluminescent Reagent (Pierce 32106) was used to fluoresce conjugates which were imaged using a BioRad Chemidock XRS+ Gel Imager.

### 1.6 PM fraction purity

To validate the separation of PM proteins from cytosolic proteins, commercially-available control antibodies were used that were validated by the manufacturer for cross-reactivity in chickens. Known PM-residing and cytosol-residing proteins targeted and their abundance was used to assess the degree of PM fractionation in flight muscle, liver, and heart samples. The membranes were incubated at 1:1000 dilution for 90 minutes at room temperature and included antibodies for 1) E-cadherin (Cell Signalling Tech. 24E10), 2) Na^+^/K^+^ ATPase (Cell Signalling Tech. 3010), 3) Glyceraldehyde-3-phosphate dehydrogenase (GAPDH) (Cell Signalling Tech. 14C10), and 4) Fatty acid translocase (FAT) (Abgent AP2883c).

### 1.7 Western Blot Band Normalisation

GLUT protein molecular weights were predicted using ExPASy (Gasteiger et al., 2005). Protein quantitation was done with a Pierce 660nm assay. 5µg of sample protein was loaded into each well of the polyacrylamide gel, in comparison with wells containing visible protein ladder (Sigma 26616). The antibody staining intensity of each Western blot sample was normalised to its corresponding total-protein stain intensity using BioRad ImageLab software. Background subtraction was applied to the total protein stain in a lane-wise fashion, while no background subtraction was applied to the antibody staining intensity. Fluorescence intensity for the total-protein stain was measured using 30% of the lane-width as per the recommendation of Gassmann et al. (2009). The antibody stain was measured using a fixed lane-width comprising of the entire lane. Normalised molecular weights were recorded.

### 1.8 Statistical Analysis

A Student’s T-test was performed for the sugar and metabolite concentrations between fed and fasted hummingbirds. We evaluated variation in isoform intensity data for each GLUT by creating linear mixed-effects models (LMMs) in R statistical language (version 3.6.1, r-project.org) using the lme4 package (Bates et al., 2015) for GLUT isoform fluorescence intensity data. We compared relative GLUT 1, 2, 3, and 5 abundance among tissues, and between fed and fasted individuals using a fully factorial design. Assumptions of residual normality were checked through visual inspection of the quantile-quantile (Q-Q) plot, a frequency histogram, and the Shapiro-Wilk Normality Test. When necessary, model parameters were transformed by a chosen function (the details of which are presented in the Results section below) resulting in the greatest homoskedasticity and data was fitted using the following formula: 

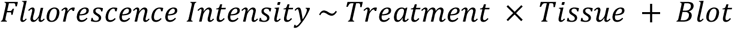

which outperformed more simplified models, as indicated by AICc (Akaike information criterion corrected for small sample sizes), the details of which are presented in Table S5. To account for the contribution of blot-to-blot variation, individual blots were treated as random effects (represented as *Blot* in the formula). Analysis of variance (ANOVA) was performed on the model parameters to determine the significance of any interactions. Post-hoc analysis was performed using the emmeans package (Lenth, 2019) within R software to determine group means and standard error. Pairwise comparison was performed to determine statistical significance of groups using the Tukey HSD method with the contrast function from the emmeans package. All data are presented as mean ± standard error.

## Results

### 2.1 Circulating Sugars and Metabolites of Fed and Fasted Hummingbirds

Overall, a significant difference was only observed for blood fructose concentrations (*t*_*9.9*_ = -17.2, *p* = 0.001) which were higher in fed hummingbirds (5.34 ± 0.2 mM) compared to fasted (0.21 ± 0.1 mM). Glucose concentrations in fed hummingbirds (30.04 ± 2.0 mM) remained similarly elevated in fasted hummingbirds (29.67 ± 1.5 mM). Lactate concentrations in fed individuals (4.31 ± 1.3 mM) were slightly lower than in fasted (6.35 ± 0.9 mM) although this was not a significant difference. Likewise, pyruvate concentrations in fed hummingbirds (0.21 ± 0.03 mM) remained elevated in fasted hummingbirds (0.22 ± 0.01 mM). These results are summarised in Figure 1.

**Figure 1:**
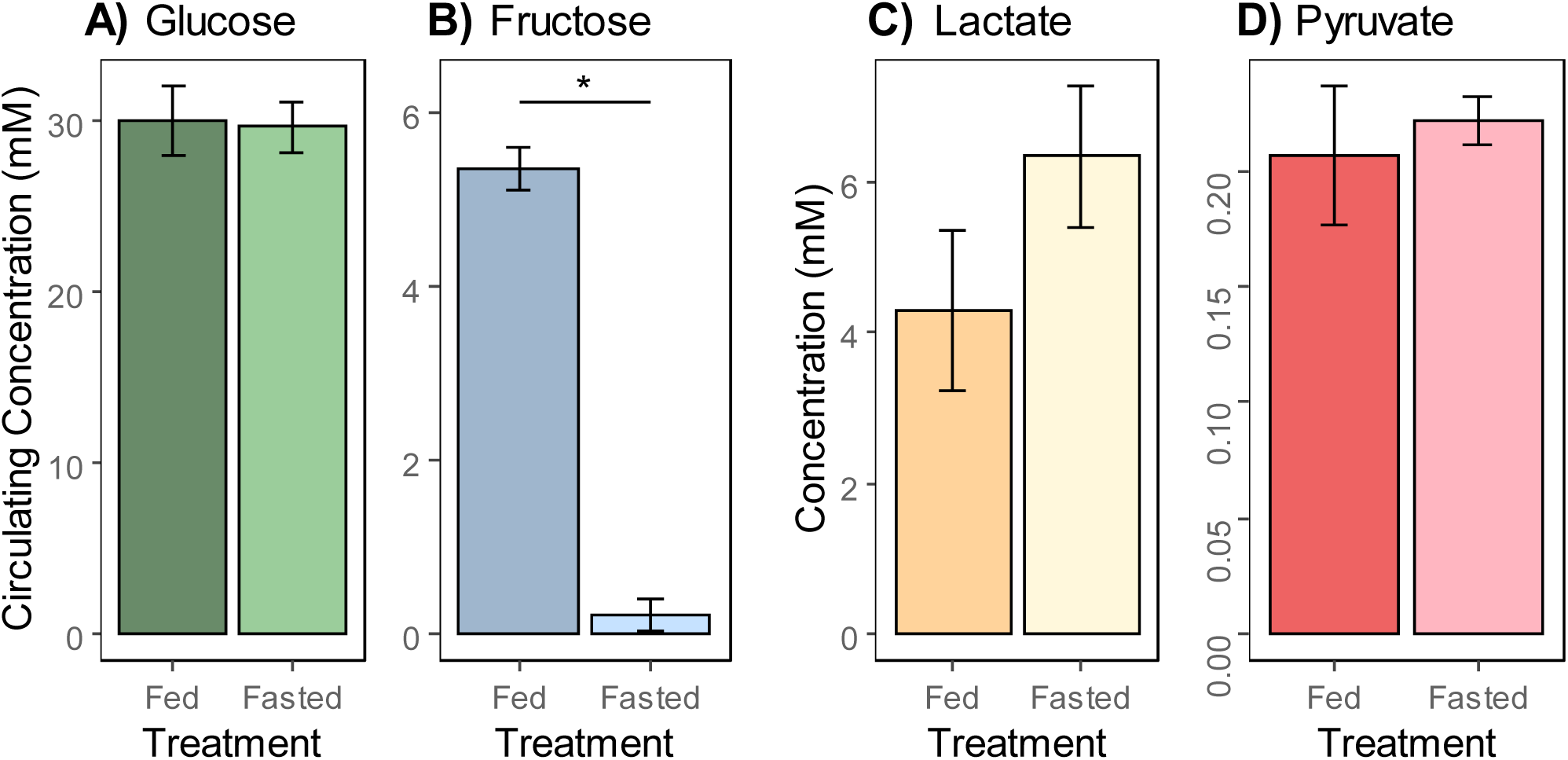
Mean concentrations (mM) ± standard error of circulating sugars A) Glucose, B) Fructose from plasma samples and metabolites C) Lactate, D) Pyruvate from whole-tissue homogenates of fed (*n = 6*) and fasted (*n = 5*). Data is presented as mean concentration in millimoles ± standard error. Asterisk (*) indicates p = 0.001.

### 2.2 Antibody Specificity and GLUT Detection

Antibodies showed a high degree of specificity for their isoform in immunoblots of HEK293 cell lysates (Table S3). In hummingbird tissue, GLUT proteins were identified by band molecular weights, and were, with one exception, present in both PM fractions and WT homogenates following PM fractionation (Table S1). GLUT1, GLUT2, GLUT3, and GLUT5 were detected in WT homogenates of flight muscle and heart tissue of ruby-throated hummingbirds, as well as in PM fractions. GLUT1 in liver WT homogenates was minimally detected and was not detected at all in liver PM fractions. GLUT1, GLUT2, and GLUT5 were detected at approximately their expected molecular weights in all tissues. GLUT3 was detected at a size slightly larger than predicted.

### 2.3 Relative GLUT Abundance

#### 2.3.1 GLUT1

With regards to the WT homogenates, no significant differences were observed in the relative abundance of GLUT1 among tissues (*F*_*2,2.5*_ = 11.58, *p* = 0.055) or the interaction of tissue and treatment (*F*_*2,13*_ = 0.262, *p* = 0.773). While WT flight muscle, regardless of treatment, had a similar GLUT1 abundance to WT heart, WT flight muscle had a significantly greater abundance compared to WT liver in both fed (flight muscle / liver ratio: 4.75 ± 1.27, *t*_*3.02*_ = 4.54, *p =* 0.040) and fasted (flight muscle / liver ratio: 5.76 ± 1.54, *t*_*3.02*_ = 4.28, *p* = 0.046) treatments. These results are summarised in Table 3 and Fig. 2A. The treatment itself, fasting, did have a significant effect (*F*_*1,13*_ *=* 7.99, *p* = 0.014) on WT GLUT1 abundance, however, multi-factor multiple comparisons using the Tukey HSD method show that only flight muscle WT GLUT1 abundance was significantly lower in fasted hummingbirds (fasted/fed ratio: 0.73 ± 0.09; *t*_*13*_ = 2.63, *p =* 0.021) (Table 1). While the effect of treatment was not significant as a whole for PM GLUT1 (*F*_*1,13.02*_ = 3.74, *p =* 0.075; Treatment), we did observe a significant effect of tissue (*F*_*1, 3.78*_ = 24, *p* = 0.009) and the interaction of tissue and treatment (*F*_*1, 13.02*_ = 17.03, *p =* 0.012). These results are summarised in Table 4 and Fig. 2B. The relative abundance of PM GLUT1 was >2-fold higher in flight muscle compared to heart within the fed treatment (Fed flight muscle / heart ratio: 4.87 ± 1.31, *t*_*4.68*_ = 5.89, *p =* 0.009). Additionally, PM GLUT1 abundance was significantly lower in flight muscle of fasted hummingbirds (fasted/fed ratio: 0.61 ± 0.06, *t*_*13*_ = 4.66, *p* = 0.002) (Table 2).

**Table 1:**
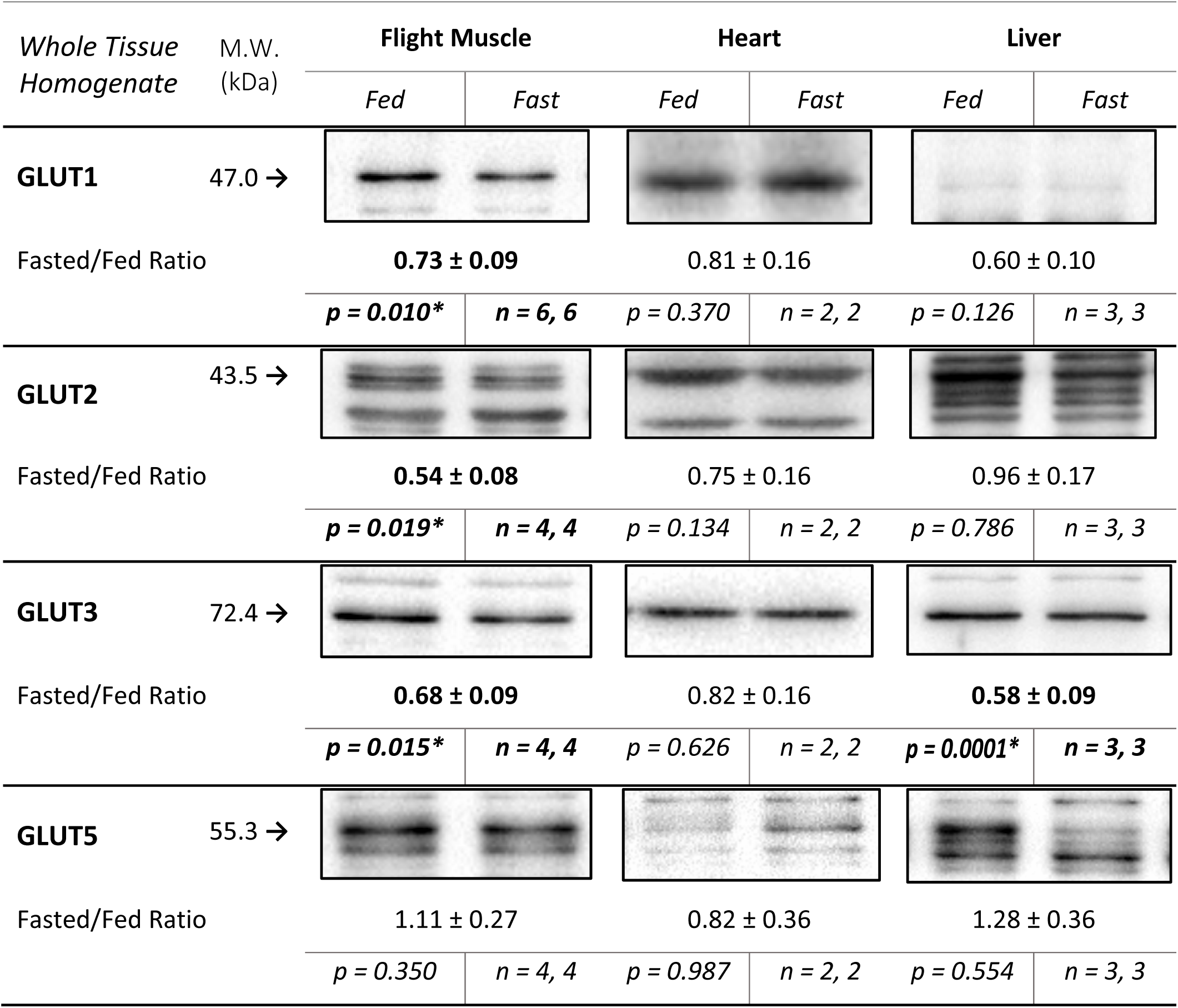
Relative WT abundance of GLUT1, GLUT2, GLUT3, and GLUT5 in flight muscle, heart, and liver of fed and fasted hummingbirds. Data and representative immunoblots are presented here for the whole tissue (WT) homogenates of hummingbird tissue. Fasted/fed ratios reflect the relative variation in GLUT protein abundance with fasting treatment. Observed molecular weights (M.W.) are reported. Sample sizes are given for the number of 1) fed hummingbirds, 2) fasted hummingbirds. Asterisks (*) indicate p < 0.05.

| Whole Tissue Homogenate | M.W. (kDa) | Flight Muscle |  | Heart |  | Liver |  |
| --- | --- | --- | --- | --- | --- | --- | --- |
|  |  | Fed | Fast | Fed | Fast | Fed | Fast |
| **GLUT1** | 47.0 → |  |  |  |  |  |  |
| Fasted/Fed Ratio |  | **0.73 ± 0.09** |  | 0.81 ± 0.16 |  | 0.60 ± 0.10 |  |
| | | $p = 0.010^*$ | $n = 6, 6$ | $p = 0.370$ | $n = 2, 2$ | $p = 0.126$ | $n = 3, 3$ |
| **GLUT2** | 43.5 → |  |  |  |  |  |  |
| Fasted/Fed Ratio |  | **0.54 ± 0.08** |  | 0.75 ± 0.16 |  | 0.96 ± 0.17 |  |
| | | $p = 0.019^*$ | $n = 4, 4$ | $p = 0.134$ | $n = 2, 2$ | $p = 0.786$ | $n = 3, 3$ |
| **GLUT3** | 72.4 → |  |  |  |  |  |  |
| Fasted/Fed Ratio |  | **0.68 ± 0.09** |  | 0.82 ± 0.16 |  | **0.58 ± 0.09** |  |
| | | $p = 0.015^*$ | $n = 4, 4$ | $p = 0.626$ | $n = 2, 2$ | $p = 0.0001^*$ | $n = 3, 3$ |
| **GLUT5** | 55.3 → |  |  |  |  |  |  |
| Fasted/Fed Ratio |  | 1.11 ± 0.27 |  | 0.82 ± 0.36 |  | 1.28 ± 0.36 |  |
| | | $p = 0.350$ | $n = 4, 4$ | $p = 0.987$ | $n = 2, 2$ | $p = 0.554$ | $n = 3, 3$ |

**Table 2:**
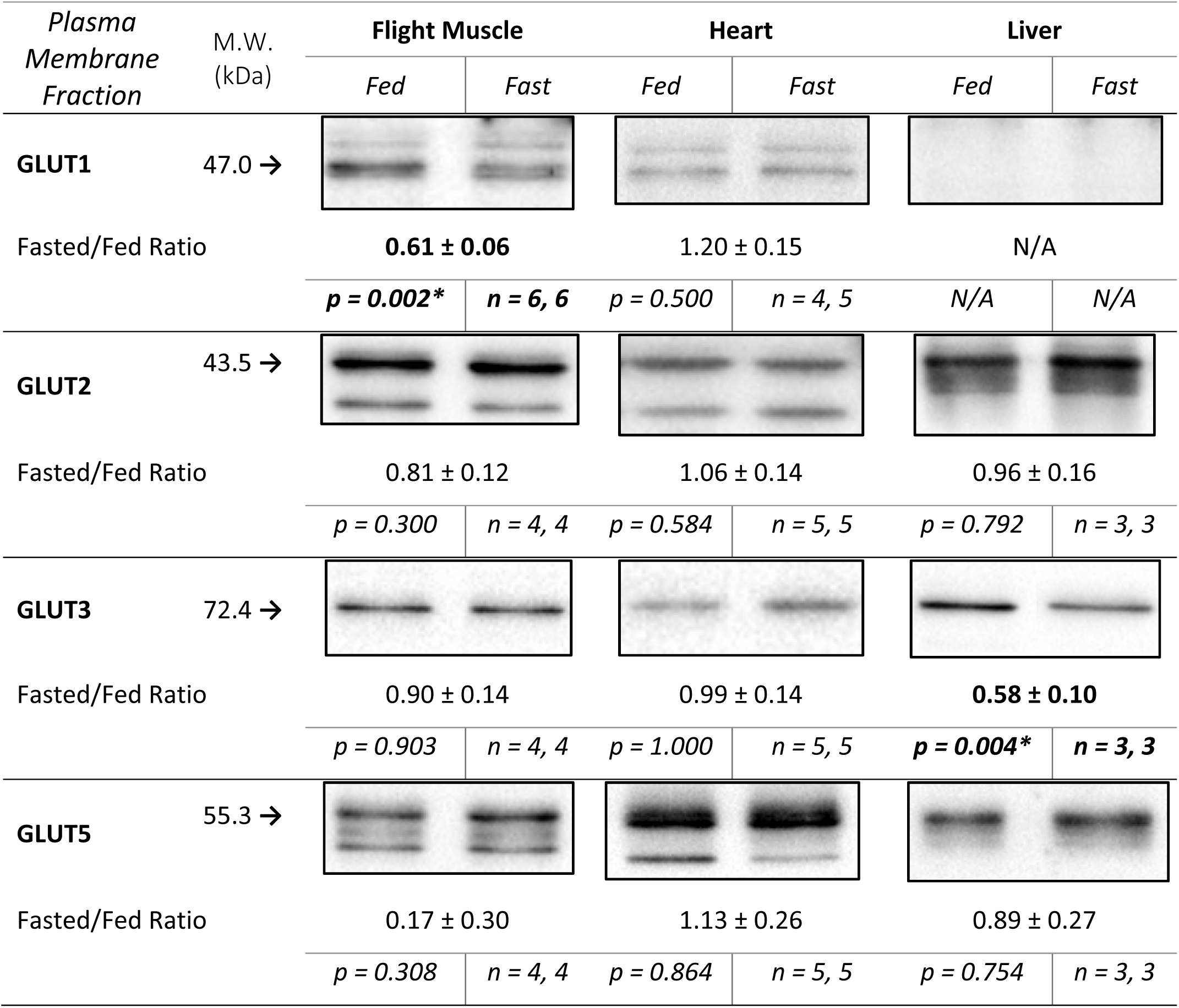
Relative PM abundance of GLUT1, GLUT2, GLUT3, and GLUT5 in flight muscle, heart, and liver of fed and fasted hummingbirds. Data and representative immunoblots are presented here for hummingbird tissue samples that underwent plasma membrane fractionation; only PM-residing GLUTs are presented. Fasted/fed ratios reflect the relative variation in GLUT protein abundance with fasting treatment. Observed molecular weights (M.W.) are reported. Sample sizes are given for the number of 1) fed hummingbirds, 2) fasted hummingbirds. Asterisks (*) indicate p < 0.05.

| Plasma Membrane Fraction | M.W. (kDa) | Flight Muscle |  | Heart |  | Liver |  |
| --- | --- | --- | --- | --- | --- | --- | --- |
|  |  | Fed | Fast | Fed | Fast | Fed | Fast |
| **GLUT1** | 47.0 → |  |  |  |  |  |  |
| Fasted/Fed Ratio | | **$0.61 \pm 0.06$** | | $1.20 \pm 0.15$ | | N/A | |
| | | $p = 0.002^*$ | $n = 6, 6$ | $p = 0.500$ | $n = 4, 5$ | N/A | N/A |
| **GLUT2** | 43.5 → |  |  |  |  |  |  |
| Fasted/Fed Ratio | | $0.81 \pm 0.12$ | | $1.06 \pm 0.14$ | | $0.96 \pm 0.16$ | |
| | | $p = 0.300$ | $n = 4, 4$ | $p = 0.584$ | $n = 5, 5$ | $p = 0.792$ | $n = 3, 3$ |
| **GLUT3** | 72.4 → |  |  |  |  |  |  |
| Fasted/Fed Ratio | | $0.90 \pm 0.14$ | | $0.99 \pm 0.14$ | | **$0.58 \pm 0.10$** | |
| | | $p = 0.903$ | $n = 4, 4$ | $p = 1.000$ | $n = 5, 5$ | $p = 0.004^*$ | $n = 3, 3$ |
| **GLUT5** | 55.3 → |  |  |  |  |  |  |
| Fasted/Fed Ratio | | $0.17 \pm 0.30$ | | $1.13 \pm 0.26$ | | $0.89 \pm 0.27$ | |
| | | $p = 0.308$ | $n = 4, 4$ | $p = 0.864$ | $n = 5, 5$ | $p = 0.754$ | $n = 3, 3$ |

**Table 3:**
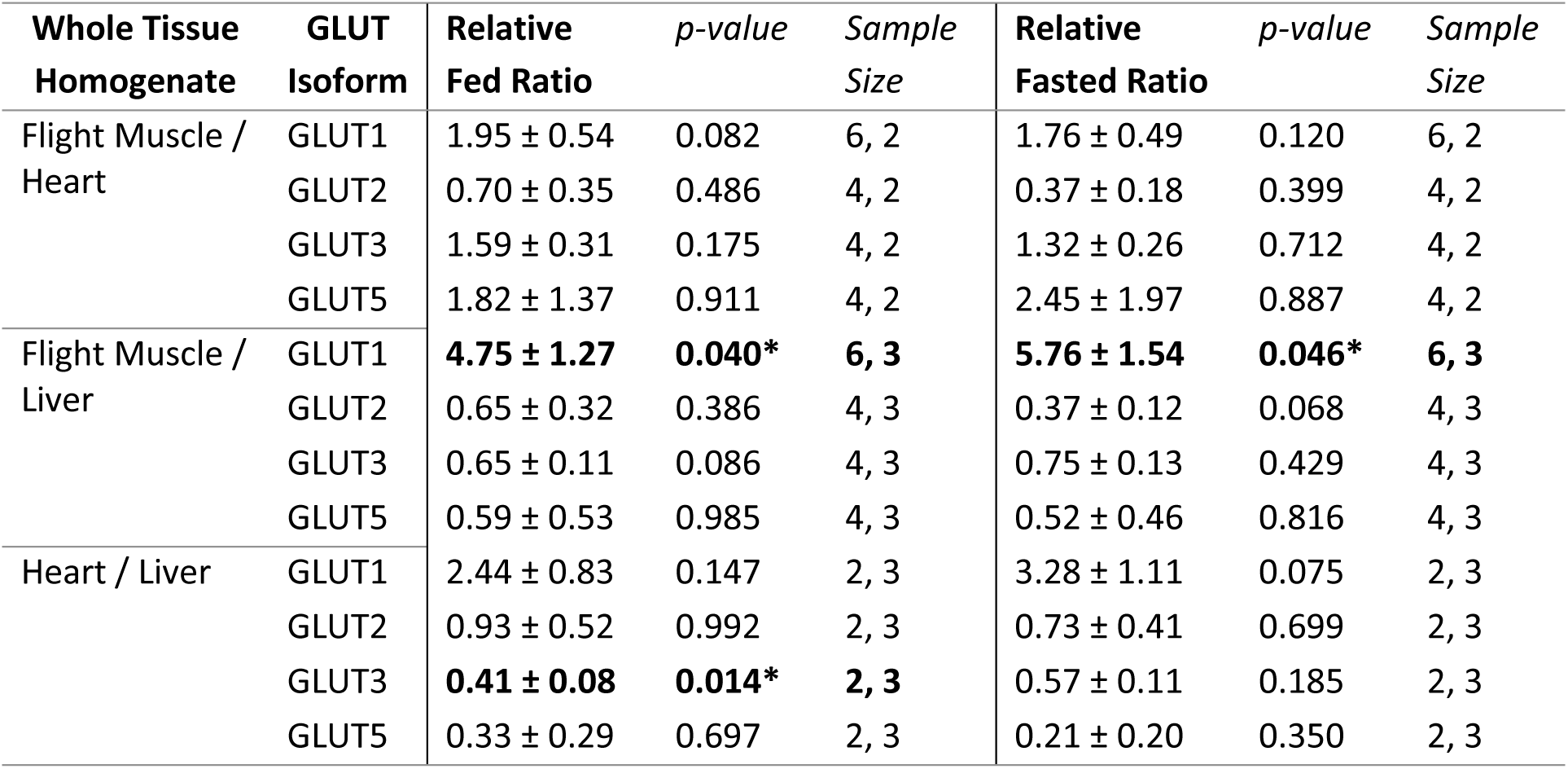
Relative abundance of GLUT1, GLUT2, GLUT3, and GLUT5 among WT homogenates compared pair-wise between flight muscle, heart, and liver of fed and fasted hummingbirds. Data represents the relative whole-tissue GLUT abundance. Asterisks (*) indicate p < 0.05.

| Whole Tissue Homogenate | GLUT Isoform | Relative Fed Ratio | *p*-value | Sample Size | Relative Fasted Ratio | *p*-value | Sample Size |
| --- | --- | --- | --- | --- | --- | --- | --- |
| Flight Muscle / Heart | GLUT1 | 1.95 ± 0.54 | 0.082 | 6, 2 | 1.76 ± 0.49 | 0.120 | 6, 2 |
|  | GLUT2 | 0.70 ± 0.35 | 0.486 | 4, 2 | 0.37 ± 0.18 | 0.399 | 4, 2 |
|  | GLUT3 | 1.59 ± 0.31 | 0.175 | 4, 2 | 1.32 ± 0.26 | 0.712 | 4, 2 |
|  | GLUT5 | 1.82 ± 1.37 | 0.911 | 4, 2 | 2.45 ± 1.97 | 0.887 | 4, 2 |
| Flight Muscle / Liver | GLUT1 | **4.75 ± 1.27** | **0.040\*** | **6, 3** | **5.76 ± 1.54** | **0.046\*** | **6, 3** |
|  | GLUT2 | 0.65 ± 0.32 | 0.386 | 4, 3 | 0.37 ± 0.12 | 0.068 | 4, 3 |
|  | GLUT3 | 0.65 ± 0.11 | 0.086 | 4, 3 | 0.75 ± 0.13 | 0.429 | 4, 3 |
|  | GLUT5 | 0.59 ± 0.53 | 0.985 | 4, 3 | 0.52 ± 0.46 | 0.816 | 4, 3 |
| Heart / Liver | GLUT1 | 2.44 ± 0.83 | 0.147 | 2, 3 | 3.28 ± 1.11 | 0.075 | 2, 3 |
|  | GLUT2 | 0.93 ± 0.52 | 0.992 | 2, 3 | 0.73 ± 0.41 | 0.699 | 2, 3 |
|  | GLUT3 | **0.41 ± 0.08** | **0.014\*** | **2, 3** | 0.57 ± 0.11 | 0.185 | 2, 3 |
|  | GLUT5 | 0.33 ± 0.29 | 0.697 | 2, 3 | 0.21 ± 0.20 | 0.350 | 2, 3 |

**Table 4:** Relative abundance of GLUT1, GLUT2, GLUT3, and GLUT5 among PM fractions of flight muscle, heart, and liver of fed and fasted hummingbirds. Values represent the relative abundance of GLUT proteins from isolated plasma membrane samples (fractionation efficiency approx. 92.1 ± 0.5%; see Table S2). Asterisks (*) indicate p < 0.05.

| Plasma Mem. Fraction | GLUT Isoform | Relative Fed Ratio | *p*-value | Sample Size | Relative Fasted Ratio | *p*-value | Sample Size |
| --- | --- | --- | --- | --- | --- | --- | --- |
| Flight Muscle / Heart | GLUT1 | **4.87 ± 1.30** | **0.009\*** | **6, 4** | 2.48 ± 0.66 | 0.075 | 6, 5 |
|  | GLUT2 | 2.31 ± 2.20 | 0.782 | 4, 5 | 1.77 ± 1.67 | 0.835 | 4, 5 |
|  | GLUT3 | 1.68 ± 0.79 | 0.814 | 4, 5 | 1.53 ± 0.71 | 0.950 | 4, 5 |
|  | GLUT5 | 1.84 ± 1.18 | 0.451 | 4, 5 | 1.96 ± 1.24 | 0.318 | 4, 5 |
| Flight Muscle / Liver | GLUT1 | *Not detected in liver PM* |  |  | *Not detected in liver PM* |  |  |
|  | GLUT2 | 1.19 ± 1.37 | 0.954 | 4, 3 | 1.00 ± 1.16 | 0.980 | 4, 3 |
|  | GLUT3 | 0.48 ± 0.27 | 0.396 | 4, 3 | 0.74 ± 0.42 | 0.958 | 4, 3 |
|  | GLUT5 | 0.61 ± 0.47 | 0.976 | 4, 3 | 0.39 ± 0.30 | 0.747 | 4, 3 |
| Heart / Liver | GLUT1 | *Not detected in liver PM* |  |  | *Not detected in liver PM* |  |  |
|  | GLUT2 | 0.51 ± 0.59 | 0.961 | 5, 3 | 0.57 ± 0.65 | 0.954 | 5, 3 |
|  | GLUT3 | 0.28 ± 0.16 | 0.225 | 5, 3 | 0.48 ± 0.27 | 0.730 | 5, 3 |
|  | GLUT5 | 0.33 ± 0.25 | 0.643 | 5, 3 | 0.20 ± 0.15 | 0.747 | 5, 3 |

**Figure 2.**
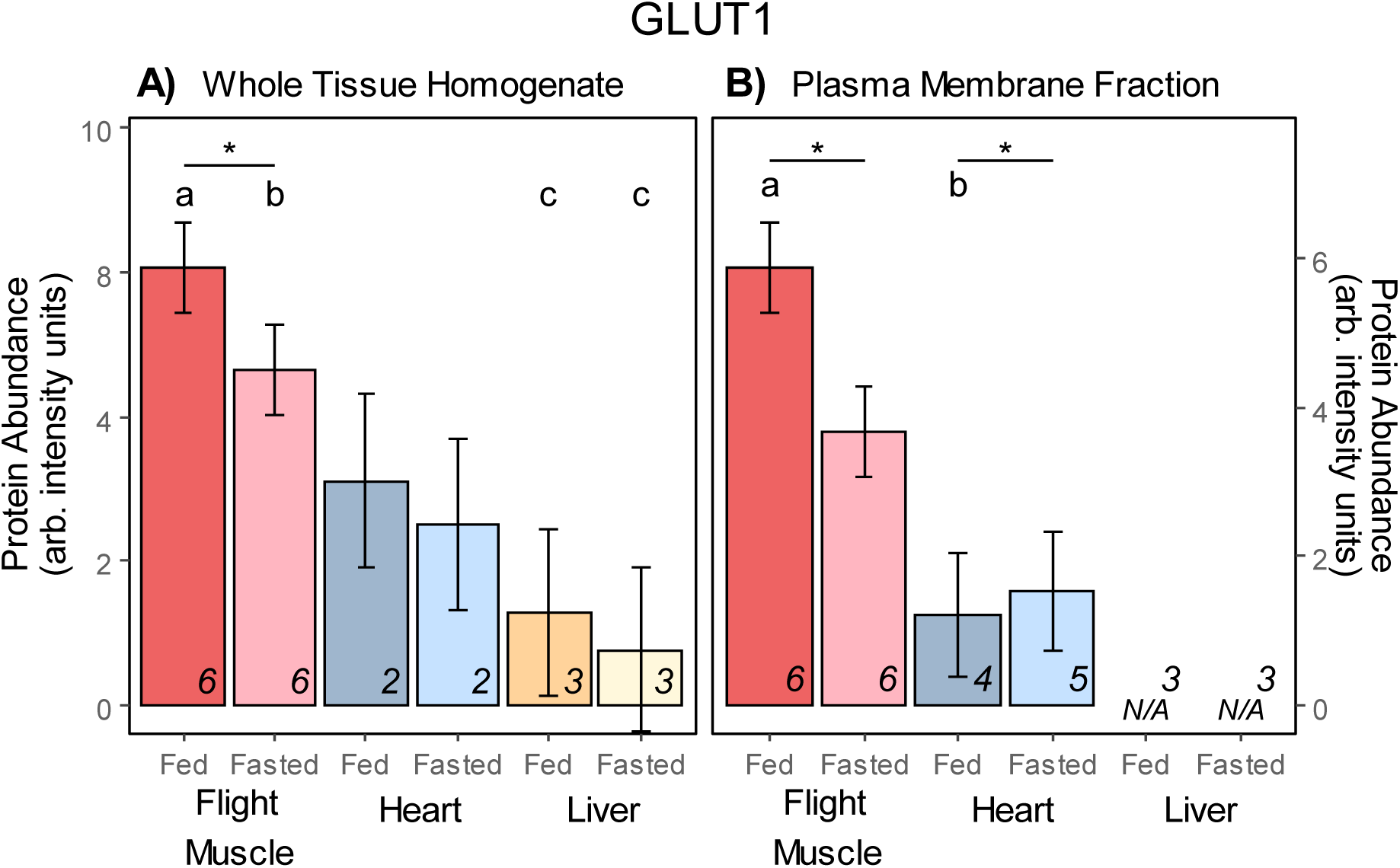
Relative protein abundance of GLUT1 in hummingbird flight muscle, heart, and liver tissue. Data represents mean ± standard error of arbitrary units of intensity based on analyses of normalised immunoblots. *Ad-libitum* fed (“Fed”) and 1-hour fasted (“Fasted”) hummingbird GLUT1 abundance was measured in A) whole tissue homogenates and B) plasma membrane fraction samples. An asterisk (*) over a tissue group indicates a significant difference (p < 0.05) of GLUT1 between fed and fasted conditions within that tissue, summarised in Table 1 and Table 2. Letters (a, b) over tissue groups represent a significant difference (p < 0.05) of GLUT1 between tissue groups in fed or fasted conditions, summarised in Table 3 and Table 4. Sample sizes are superimposed on the bottom-right for each tissue and treatment.

#### 2.3.2 GLUT2

Amongst WT homogenates, a significant effect of treatment was observed regarding WT GLUT2 relative abundance (*F*_*1, 11*_ = 6.22, *p =* 0.029). Multiple comparisons revealed that only flight muscle had a significantly lower WT GLUT2 abundance in fasted hummingbirds (fasted/fed ratio: 0.54 ± 0.08, *t*_*14.5*_ = 2.63, *p =* 0.019), while heart and liver tissue did not show a significant difference (Table 1 and Fig. 3A). Regardless of feeding or fasting treatment, no significant difference was observed in WT GLUT2 relative abundance among tissues (Table 3). No significant difference in relative abundance was noted for tissue or the interaction of tissue and treatment for WT GLUT2. Further, no significant difference was observed among tissues, treatment, or the interaction of tissue and treatment for PM GLUT2 (Table 2, Table 4, and Fig. 3B).

**Figure 3.**
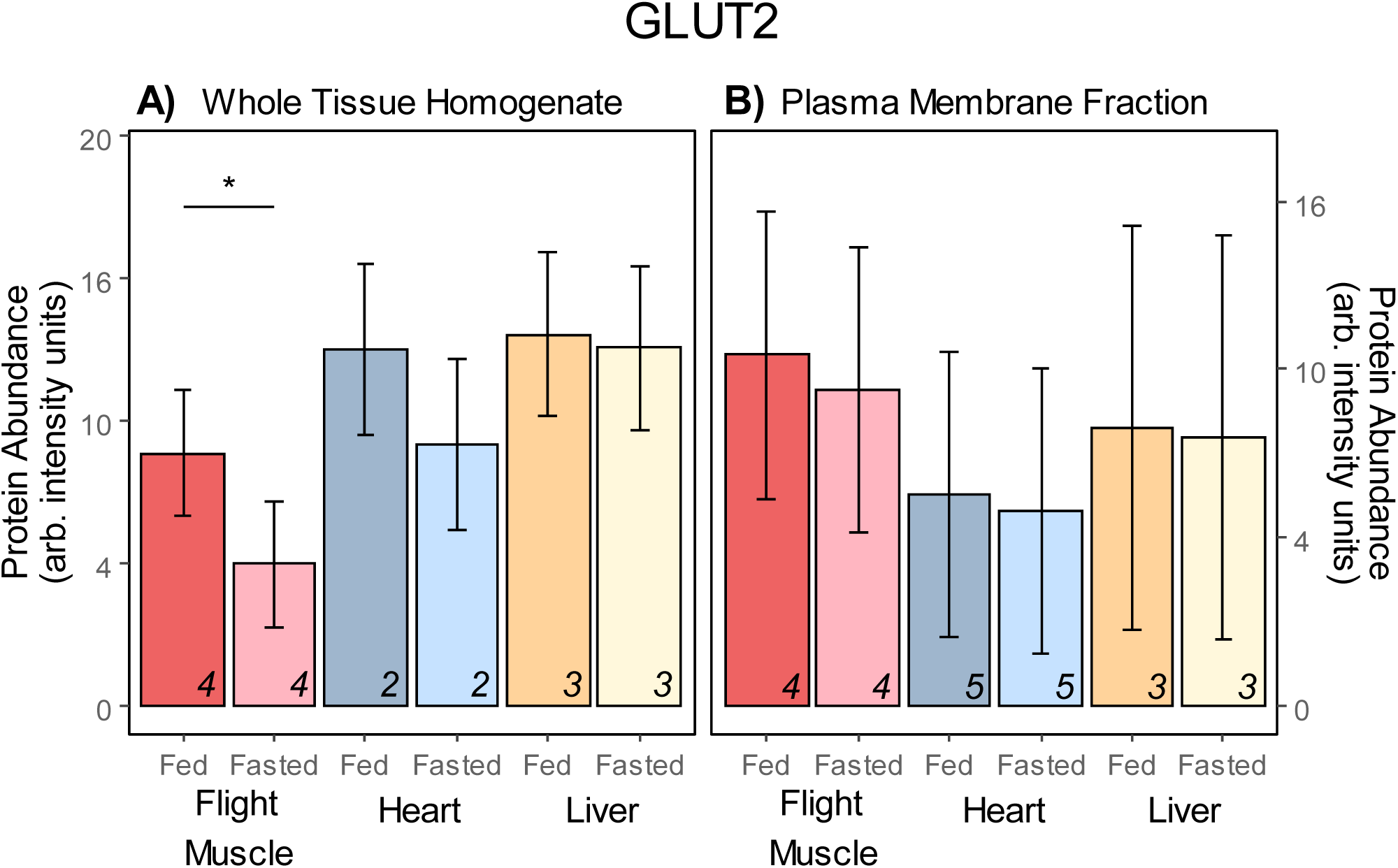
Relative protein abundance of GLUT2 in hummingbird flight muscle, heart, and liver tissue. Data represents mean ± standard error of arbitrary units of intensity based on analyses of normalised immunoblots. *Ad-libitum* fed (“Fed”) and 1-hour fasted (“Fasted”) hummingbird GLUT2 abundance was measured in A) whole tissue homogenates and B) plasma membrane fraction samples. An asterisk (*) over a tissue group indicates a significant difference (p < 0.05) of GLUT2 between fed and fasted conditions within that tissue, summarised in Table 1 and Table 2. Differences in abundance of GLUT2 between tissue groups in fed or fasted conditions, summarised in Table 3 and Table 4. Sample sizes are superimposed on the bottom-right for each tissue and treatment.

#### 2.3.3 GLUT3

Fasting significantly affected the relative abundance of WT GLUT3 (*F*_*1, 11*_ = 17.08, *p =* 0.002). Multi-factor multiple comparisons with the Tukey HSD method revealed that both flight muscle (fasted/fed ratio: 0.68 ± 0.09, *t*_*24.8*_ = 2.61, *p =* 0.015) and liver (fasted/fed ratio: 0.58 ± 0.09, *t*_*24.8*_ = 4.58, *p =* 0.0001) had significantly less WT GLUT3 in fasted hummingbirds, while no significant difference was observed in heart WT homogenates (Table 1 and Fig. 4A). No significant difference was observed for relative WT GLUT3 abundance among tissues or the interaction of tissue and treatment. Regardless of treatment, WT GLUT3 abundance was similar among tissues with the exception of the liver having significantly greater relative WT GLUT3 abundance compared to heart in fed hummingbirds (fed liver / heart ratio: 2.46 ± 0.46, *t*_*3.5*_ = 5.83, *p =* 0.014) (Table 3 and Fig. 4B). In PM fractions, a significant effect of the fasting treatment was observed on the relative PM GLUT3 abundance (*F*_*1,16*_ = 13.13, *p =* 0.002). No significant difference was observed among tissues (Table 4), however, the interaction of tissue and treatment was significant (*F*_*2,16*_ = 6.46, *p =* 0.009). Through multiple comparisons, it was observed that only liver PM GLUT3 relative abundance was significantly lower in fasted hummingbirds (fasted/fed ratio: 0.58 ± 0.14, *t*_*16*_ *=* 4.54, *p =* 0.004) (Table 2).

**Figure 4.**
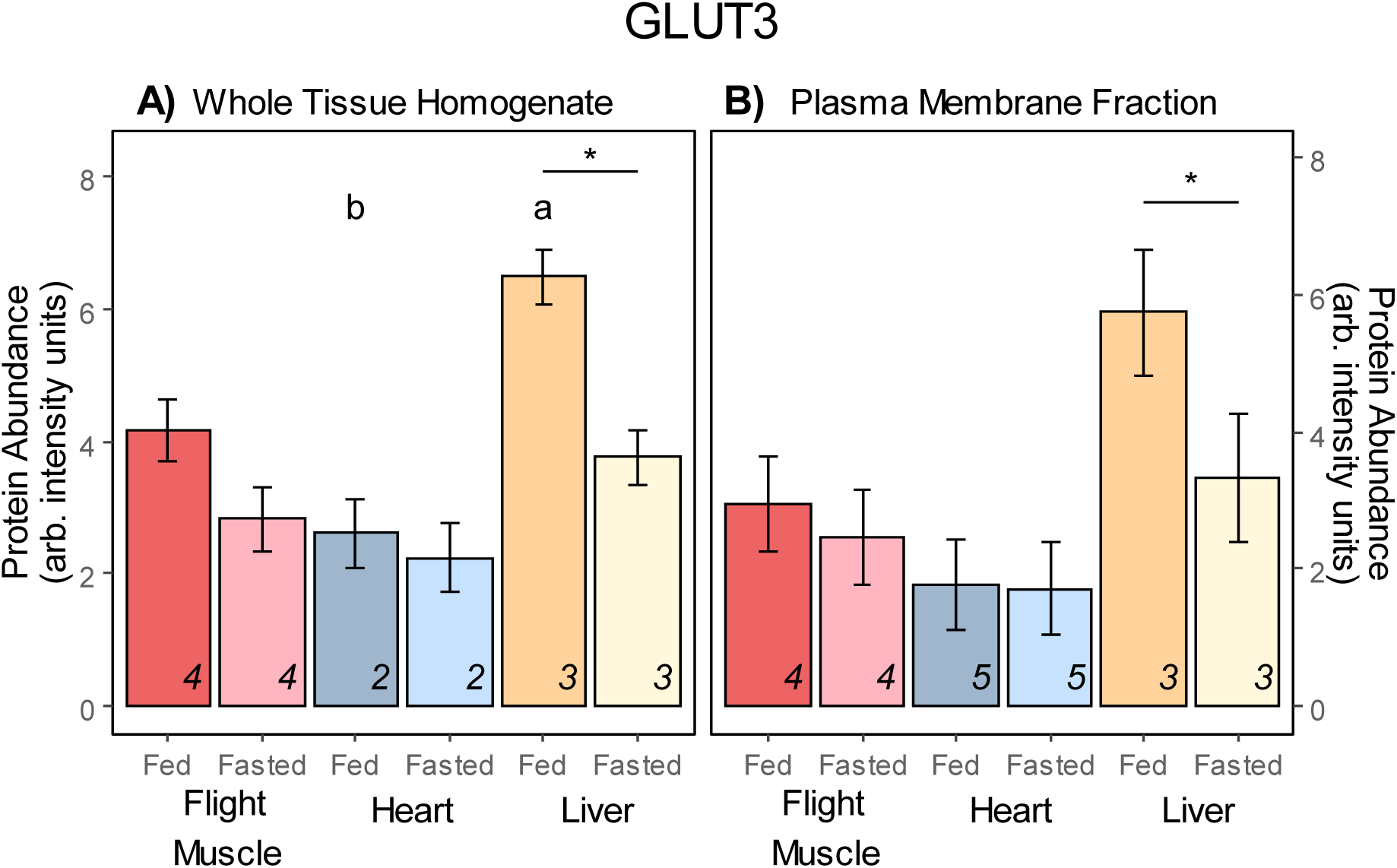
Relative protein abundance of GLUT3 in hummingbird flight muscle, heart, and liver tissue. Data represents mean ± standard error of arbitrary units of intensity based on analyses of normalised immunoblots. *Ad-libitum* fed (“Fed”) and 1-hour fasted (“Fasted”) hummingbird GLUT3 abundance was measured in A) whole tissue homogenates and B) plasma membrane fraction samples. An asterisk (*) over a tissue group indicates a significant difference (p < 0.05) of GLUT3 between fed and fasted conditions within that tissue, summarised in Table 1 and Table 2. Letters (a, b) over tissue groups represent a significant difference (p < 0.05) of GLUT3 between tissue groups in fed or fasted conditions, summarised in Table 3 and Table 4. Sample sizes are superimposed on the bottom-right for each tissue and treatment.

#### 2.3.4 GLUT5

No significant effect of tissue or treatment, or their interaction, were observed for the relative abundance of WT GLUT5. Regardless of treatment WT GLUT5 relative abundance did not differ significantly between tissues (Table 1, Table 3, Fig. 5A). PM GLUT5 did not show any significant effect with tissue, treatment, or their interaction. No significant effect was observed in any tissue with fasting treatment (Table 2, Fig. 5B). Regardless of feeding or fasting, no significant difference was observed in the relative PM GLUT5 abundance among tissues (Table 4).

**Figure 5.**
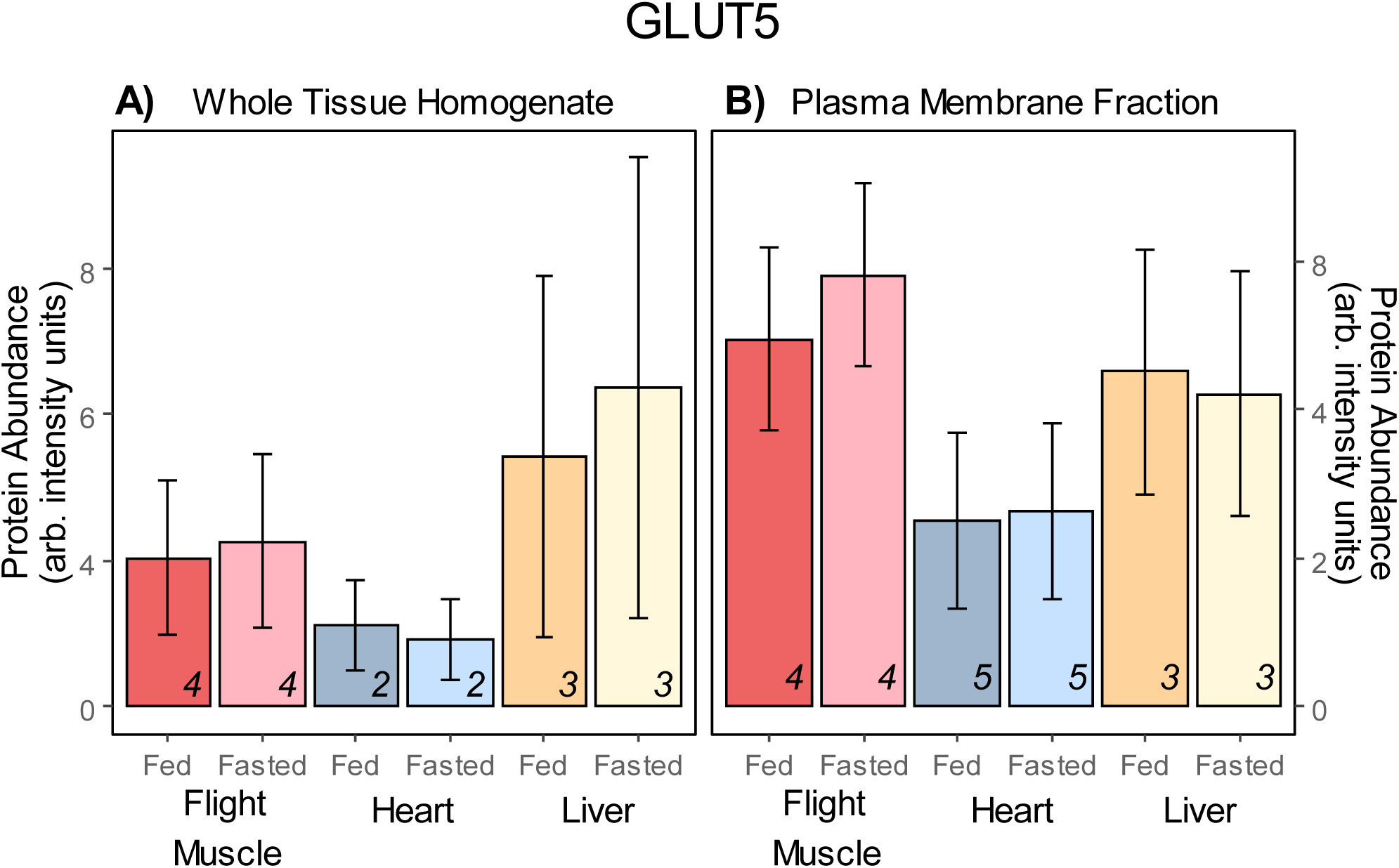
Relative protein abundance of GLUT5 in hummingbird flight muscle, heart, and liver tissue. Data represents mean ± standard error of arbitrary units of intensity based on analyses of normalised immunoblots. *Ad-libitum* fed (“Fed”) and 1-hour fasted (“Fasted”) hummingbird GLUT5 abundance was measured in A) whole tissue homogenates and B) plasma membrane fraction samples. Differences in GLUT5 abundance between fed and fasted conditions within a given tissue are summarised in Table 1 and Table 2. Differences in overall GLUT5 abundance between tissue groups in fed or fasted conditions, summarised in Table 3 and Table 4. Sample sizes are superimposed on the bottom-right for each tissue and treatment.

## Discussion

Following a 1-hour treatment period, hummingbirds that were fasted (*n* = 5) had significantly lower blood fructose concentration compared to those that continued to feed (*n* = 6) (fed; 5.34 ± 0.24 mM, fasted; 0.21 ± 0.15 mM, *t*_*9.9*_ = -17.2, *p* > 0.001; Figure 1). As this is the first report of blood fructose concentrations in hummingbirds, it is useful to compare our results against available data from other vertebrates that specialise on sugar-rich food sources. In frugivorous bats, such as the Egyptian fruit bat (*Rousettus aegyptiacus*), blood fructose concentrations are known to rise to ∼11mM following a fructose-only meal (Keegan, 1977). Egyptian fruit bats, much like hummingbirds, have been shown to rapidly incorporate fructose into their pool of metabolizable substrates (Keegan, 1977). In the nectarivorous Pallas’s long-tongued bat (*Glossophaga soricina*), the fraction of expired CO_2_ supported by labelled carbons (*f*_*exo*_) from a fructose meal takes ∼9 minutes to reach 50% (Voigt and Speakman, 2007) while it took ruby-throated hummingbirds ∼14 minutes (Chen and Welch, 2014). In this study, we also see very low blood fructose concentrations in fasted hummingbirds compared to those that were fed (Figure 1). We further observed a slightly higher lactate concentration in fasted hummingbirds, although not significantly so (Figure 1), suggesting elevated fructolytic pathway activity (Dekker et al., 2010). These results indicate a rapid depletion of circulating fructose levels and may imply the rapid incorporation of exogenous blood fructose into the pool of metabolizable substrates in hummingbirds entering a fast.

In contrast, while circulating concentrations of glucose were, as expected, high in fed hummingbirds, they remained elevated in fasted hummingbirds (fed; 30.04 ± 2.03 mM, fasted; 29.67 ± 1.25 mM; Fig. 1). Beuchat and Chong (1998) had previously observed a similar trend in hummingbirds entering a fast; blood glucose concentration remained elevated for the first hour and declined only after ∼1.5 hours of fasting. Organs such as the brain are exceptionally demanding of glucose (Tokushima et al., 2005) in *gallus gallus* chicks and likely other birds as well. Further, lipogenic pathways of the hummingbird liver also shows a preference for glucose over fructose (Dick et al., 2019). Finally, while hummingbirds have the capacity to hover oxidising either glucose-only or fructose-only meals (Chen and Welch, 2014), their flight muscle cells’ maximal capacity for monosaccharide phosphorylation is twice as high for glucose compared to fructose in tissue homogenates *in vitro* (Myrka and Welch, 2018). As hummingbird muscles lack extensive glycogen stores (Suarez et al., 1990), processes such as gluconeogenesis in the liver or other tissues may underlie the maintenance of elevated blood glucose. Our observations suggest that glucose uptake capacity is initially downregulated in hummingbirds entering a fast while fructose uptake capacity is unchanged.

Control of glucose and fructose flux is well-described in avian species. Despite the absence of the insulin-GLUT4 system in avian muscle cells (Dupont, 2009), chickens and English sparrows (*Passer domesticus*) have demonstrated coordinated expression of GLUT isoforms to control sugar transmembrane transport (Sweazea and Braun, 2006; Wagstaff and White, 1995). Less is known about hummingbird GLUT expression and regulation. In this study, we detected a strong immunoblot signal of the protein presence of GLUT 2, 3 and 5 in hummingbird flight muscle, heart, and liver tissue in WT homogenates (Table 1). GLUT2 was observed as a doublet while GLUT3 was detected at a size slightly larger than predicted, both of which may be attributable to variations in glycosylation (Asano et al., 1992; Ohtsubo et al., 2013). GLUT1 protein was detected in hummingbird flight muscle and heart (Table S1). GLUT1 protein in WT liver homogenates of ruby-throated hummingbirds was only minimally visible (Table 1) and was, surprisingly, not detected in PM fractions (Table 2). This result is in contrast to previously reported detection of hepatic mRNA transcript for GLUT1 in both chickens (Byers et al., 2018) and hummingbirds (Welch et al., 2013). However, as GLUT1 is abundant in erythrocytes (Carruthers, 2009), it is possible that the previous mRNA detection, as well as our detection of some hepatic GLUT1 protein, may have resulted from red blood cell contamination. While the presence of transcript does not necessarily mean that the final protein form is being fully transcribed (Vogel and Marcotte, 2012), it is clear that hepatic GLUT1 is not translocated to the plasma membrane. Our findings are similar to others that have failed to detect GLUT1 in the avian liver (Byers et al., 2017; Carver et al., 2001), raising the possibility that the role of hepatic GLUT1 protein may be much more reduced among birds than previously appreciated.

In chickens, GLUT protein expression appears to be dependent on synthesis or degradation of protein (Yamada et al., 1983) rather than the translocation from cytosolic pools that is observed in mammalian cells (Guma et al., 1995). If the same were true in hummingbirds, GLUT abundance of the overall tissue should be tied to the abundance of GLUT protein in the PM. In this study, we noted that flight muscle overall showed the greatest response to fasting, in terms of relative WT GLUT abundance. We detected significantly lower WT GLUT1 (fasted/fed ratio: 0.73 ± 0.09, *p* = 0.010, *N fed:fasted =* 6, 6; Table 1 and Fig. 2A), WT GLUT2 (fasted/fed ratio: 0.54 ± 0.08, *p* = 0.019, *N fed:fasted =* 4, 4; Table 1 and Fig. 3A), and WT GLUT3 (fasted/fed ratio: 0.68 ± 0.09, *p* = 0.015, *N fed:fasted =* 4, 4; Table 1 and Fig. 4A) in flight muscle of fasted hummingbirds. While GLUTs do not contribute to transmembrane transport of sugars until they are expressed in the PM, this reduction of glucose-specific WT GLUTs across the whole flight muscle tissue may underlie the reduced glucose uptake capacity. This may be especially important in their flight muscle as its metabolic demands overshadow that of other tissues during hovering (Suarez, 1992). Heart tissue of fasted hummingbirds showed no differences in GLUT abundance compared to fed hummingbirds. This muted response to fasting was expected as cardiac metabolism relies predominantly on circulating triglycerides (Pascual and Coleman, 2016) and this may be especially true of hummingbirds as they routinely switch to fatty acid metabolism during periods of fasting (Welch et al., 2018). However, it may also imply that the elevated blood glucose concentration in fasted hummingbirds provides sufficient substrate for cardiac metabolism, especially given hummingbirds used in this study were constrained to continuously perch during the fasting period. Finally, in liver tissue, only WT GLUT3 was significantly lower in fasted hummingbirds liver (fasted/fed ratio: 0.58 ± 0.09, *p* = 0.0001, *N fed:fasted =* 3, 3; Table 1 and Fig. 4A). Chickens have also been shown to decrease their hepatic rate of glucose metabolism when fasted (Goodridge, 1968). And considering that we did not detect PM GLUT1 protein, as described previously, this reduction in GLUT3 abundance during a fast in the liver might have a large effect on glucose import capacity.

Despite the relative abundance in WT homogenates, the functional capacity for sugar import into an active cell is dependent on the density of active GLUTs expressed in the PM (Wasserman, 2009). In this study, we detected significantly less PM GLUT1 protein in the flight muscle (fasted/fed: 0.61 ± 0.06, *p* = 0.002, *N fed:fasted =* 6, 6; Table 2 and Fig. 2B) and PM GLUT3 protein in the liver (fasted/fed ratio: 0.58 ± 0.10, *p* = 0.004, *N fed:fasted =* 3, 3; Table 2 and Fig 4B) of fasted hummingbirds. This study is the first to report differences in subcellular abundance of GLUT protein in fed and fasted hummingbirds. Our results suggest that within the first hour of a fast, hummingbirds maintain elevated blood glucose levels through the lowering of glucose-specific glucose transporter abundance in the PM of these tissues. In this case, reduced expression of two high-affinity glucose-specific GLUTs in the PM, GLUT1 (*K*_*m*_ ≈ 3-5mM; Zhao & Keating, 2007) and GLUT3 (*K*_*m*_ ≈ 1.5mM; Mueckler & Thorens, 2013), may substantially impact the import of glucose into flight muscle and liver tissues, respectively. As we observed concordant decreases in WT GLUT1 in the flight muscle and WT GLUT3 in the liver, our data suggests that hummingbirds, much like chickens, regulate PM GLUT expression via synthesis or degradation of protein, rather than its translocation. Additionally, a recent study measuring levels of chicken GLUT1 mRNA also noticed a decrease in transcript following fasting (Coudert et al., 2018). We further observed that the fructose-transporting GLUT2 (Fig. 3B) and GLUT5 (Fig. 5B), did not change in PM abundance in any tissues tested following the 1-hour fast (Table 2). GLUT5 abundance did not change in WT homogenates either for any tissues. This suggests that PM GLUT5 and PM GLUT2 remain constitutively expressed in the PM of hummingbirds entering a fast. As expression of PM GLUTs allows for rapid sugar import (Wasserman, 2009), and as the highest affinity for fructose that is exhibited by GLUT5 (*K*_*m*_ ≈ 11-12mM; Douard & Ferraris, 2008), this constitutive expression may underlie the observed reduced blood fructose concentration in fasted hummingbirds.

In conclusion, we detected GLUTs 1, 2, 3, and 5 in all tissues, with the exception of GLUT1 in the liver PM. Flight muscle was observed to respond most dynamically to a 1-hour fast, followed by the liver, and finally the heart. We observed a decrease in the PM and WT abundance of glucose-specific GLUT1 in flight muscle and GLUT3 in the liver, which may lead to reduced glucose import capacity and thus maintenance of elevated blood glucose concentrations in fasted hummingbirds. In addition, we observed the constitutive expression of fructose-transporting PM GLUT2 and PM GLUT5 in all tissues, which should permit continued fructose uptake into theses tissue during initial stages of fasting, leading to near-depletion of the circulating pool of fructose. We further observed that the changes in GLUT protein expression occur both intracellularly and in the PM – no decrease of GLUT protein in the PM occurred without a concordant decrease in WT homogenates. These results suggest that hummingbirds, similar to other birds, may rely on mechanisms of GLUT synthesis and degradation, rather than translocation, to regulate extreme fluxes in circulating glucose and fructose concentrations.

## List of Abbreviations

AIC: Akaike Information Criterion
AICc: Akaike Information Criterion for small sample sizes
ANOVA: Analysis of Variance
APS: Ammonium Persulfate
cDNA: Complementary Deoxyribonucleic Acid
CO_2_: Carbon Dioxide
DTT: Dithiothreitol
*f*_exo_: Proportion of expired CO_2_ fuelled by oxidation of exogenous sugar
GAPDH: Glyceraldehyde-3-Phosphate Dehydrogenase
GLUT: Glucose Transporter
HEK293T: Homo sapiens Embryonic Kidney cell line with Mutant SV40 large T antigen
HRP: Horseradish Peroxidase
LC MRM/MS: Liquid Chromatography Multiple Reaction Monitoring Mass Spectrometry
LMM: Linear Mixed-effects Model
mM: Millimolar
mRNA: Messenger Ribonucleic Acid
MW: Molecular Weight
NCBI: National Center for Biotechnology Information
NP-40: Nonidet P-40
PBST: Phosphate Buffered Saline with Tween 20
PM: Plasma Membrane
PVDF: Polyvinylidene Fluoride
QQ plot: Quantile-quantile plot
RIPA: Radioimmunoprecipitation Assay
SDS: Sodium Dodecyl Sulfate
SDS PAGE: Sodium Dodecyl Sulfate Polyacrylamide Gel Electrophoresis
TEMED: Tetramethylethylenediamine
TMIC: The Metabolomics Innovation Centre
UTSC: University of Toronto Scarborough Campus
WT: Whole Tissue

## Acknowledgements

We would like to thank Alex Myrka for his lessons on Western blotting and Lily Hou for her expertise in the development of the mixed-effects model employed in this study. We are also grateful to the Welch Lab volunteers for their assistance with hummingbird capture. We extend a special thanks to Rachael Sparklin and Dr. Winston Timp for assembling the hummingbird transcriptome. Finally, we would like to thank Dr. Aarthi Ashok, Dr. Mauricio Terebiznik, and the Welch Lab team for their insights, critiques, and support.

## Supplementary Materials

### 3.1 Custom antibodies: GLUT1, 2, 3, and 5 detection in PM and WT homogenates

**Table S1:**
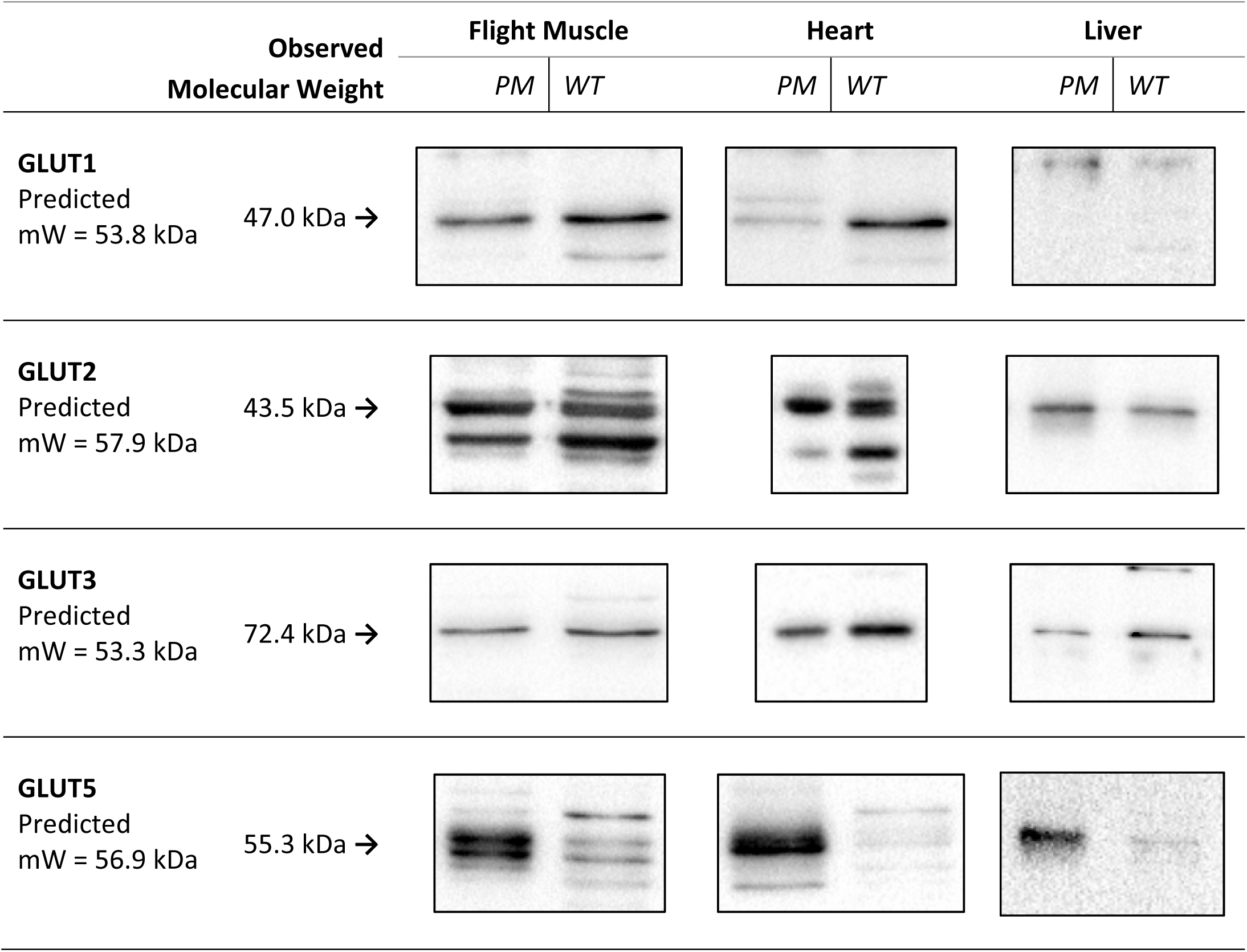
GLUTs 1, 2, 3, and 5 observed molecular weights in plasma membrane (PM) fractions and whole-tissue (WT) homogenates of flight muscle, heart, and liver. Representative immunoblots are shown for each tissue and fraction.

### 3.2 Plasma Membrane Fractionation Purity

**Table S2:**
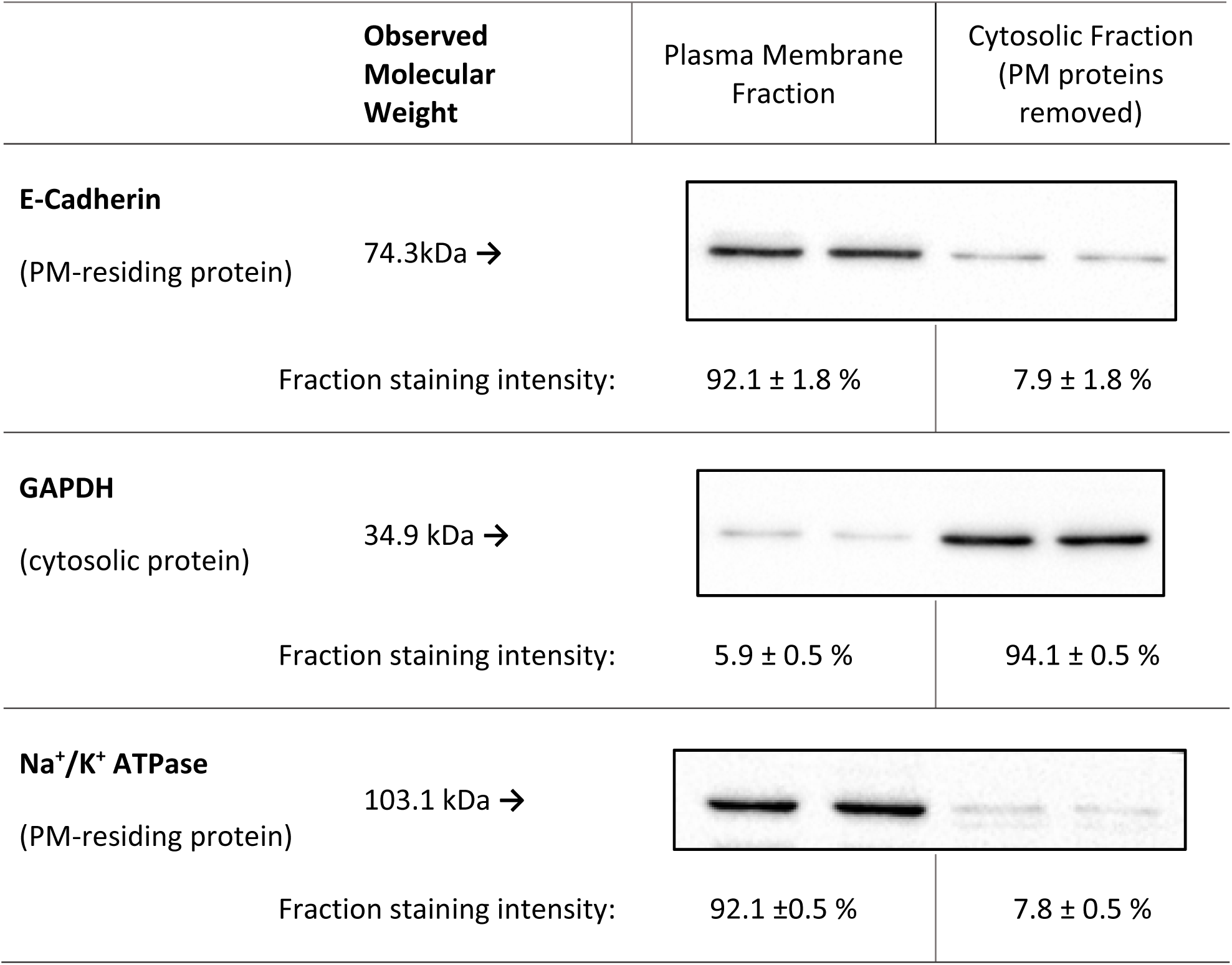
Relative distribution of known cysotolic or PM-residing proteins following PM fractionation. Fraction purity indicates the relative abundance of protein in either the PM-only fraction compared to the without-PM-fraction (i.e. cytosolic proteins only).

### 3.3 GLUT Amino Acid Sequence and Antibody Epitope

**Table S3:**
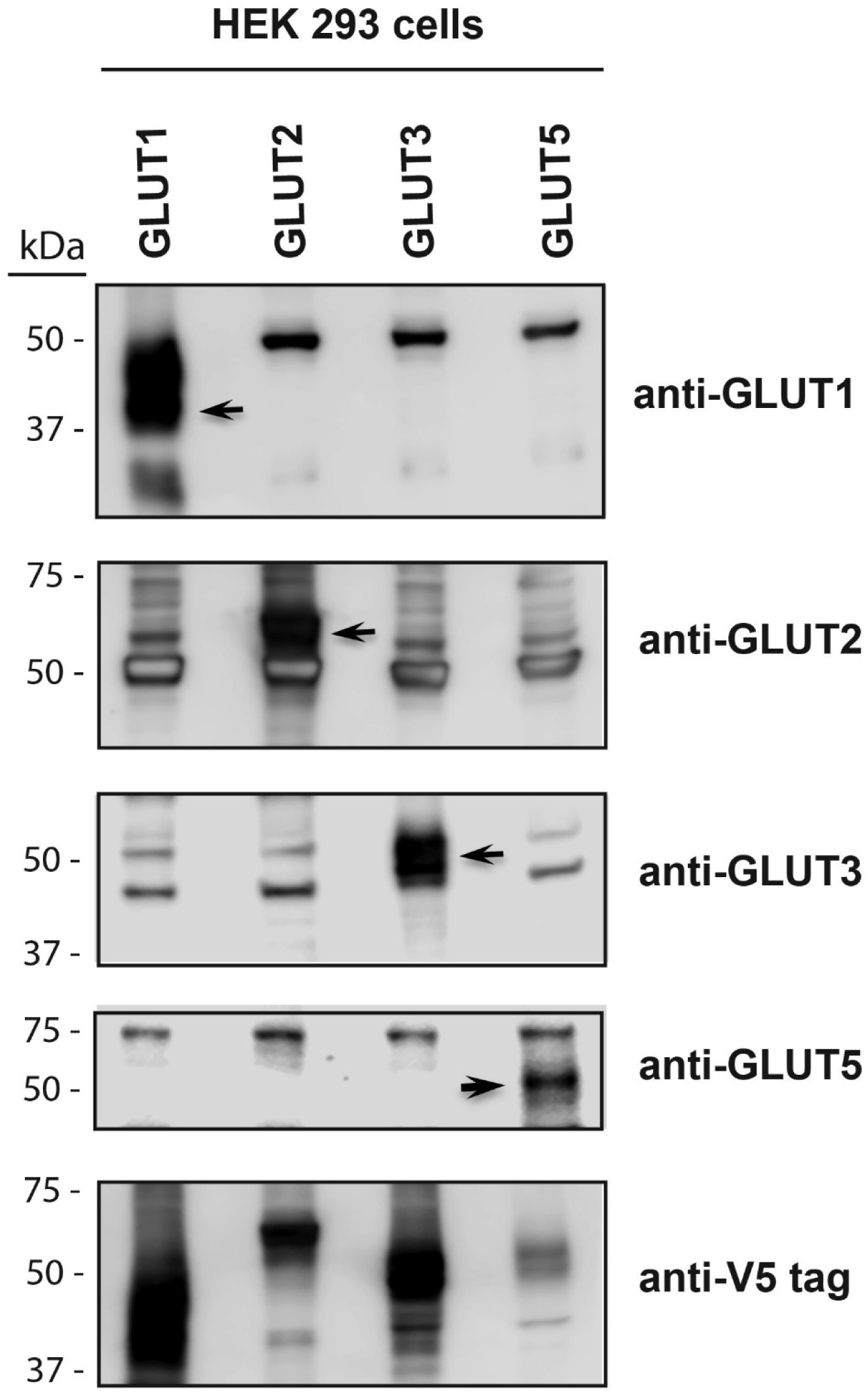
Immunoblots on lysates of overexpressed GLUT1, GLUT2, GLUT3, GLUT5 protein. Each immunoblot lane represents a cell lysate produced from an entire well of a 6-well cell-culture dish. Isoform specificity was tested via immunoblotting all cell lysates (empty vector control, acGLUT1, 2, 3, and 5) with each novel GLUT antibody and observing GLUT protein signal overlap.

**Table S4:**
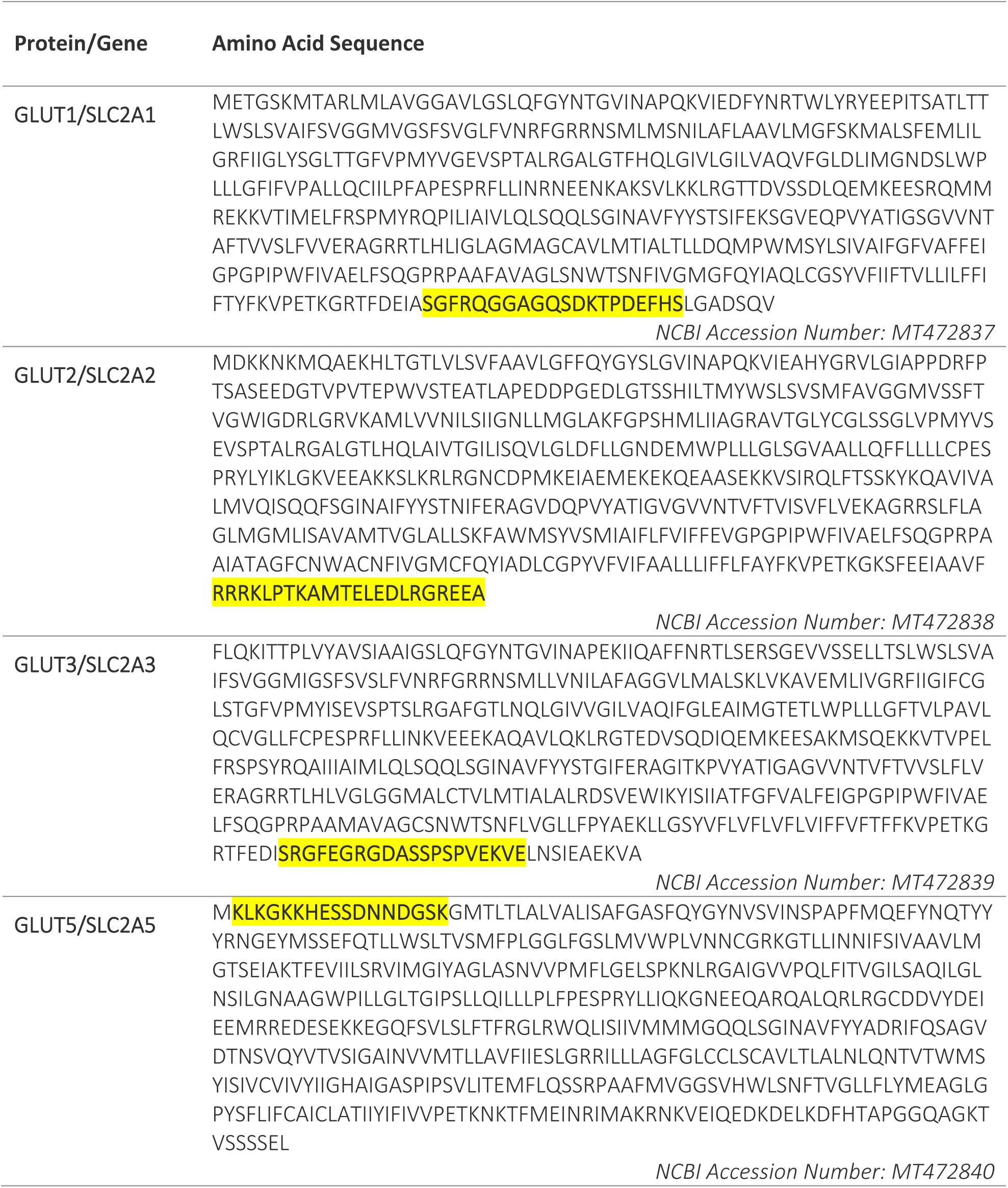
Ruby-throated hummingbird specific GLUT1, GLUT2, GLUT3 and GLUT5 protein sequences. Highlighted regions indicated epitope targeted during antibody development to ensure greatest dissimilarity between targeted isoforms

### 3.4 AIC Scores

**Table S5:** Akaike information criterion (AIC) and AIC with corrections for small sample size (AICc) scores presented for each GLUT isoform model. Due to a relatively small sample size, AICc was preferred over AIC. Models with the lowest AICc score were selected for post hoc analysis and are indicated with an asterisk (*). The models tested are as follows: 1: ***Fluorescence Intensity*** ∼ ***Treatment*** + ***Blot*** 2: ***Fluorescence Intensity*** ∼ ***Tissue*** + ***Blot*** 3: *Fluorescence Intensity* ∼ *Treatment* + *Tissue* + *Blot* 4: *Fluorescence Intensity* ∼ *Treatment* × *Tissue* + *Blot*

| GLUT | Model | AIC Score | AICc Score | AIC Score | AICc Score |
| --- | --- | --- | --- | --- | --- |
|  |  | WT | WT | PM | PM |
| **GLUT1** | 1 | 515 | 519 | 31.9 | 34.4 |
|  | 2 | 516 | 519 | 23.9 | 26.4 |
|  | 3 | 479 | 485 | 25.2 | 29.2 |
|  | **4** | **446** | **455\*** | **17.4** | **23.4\*** |
| **GLUT2** | 1 | 604 | 607 | 812 | 814 |
|  | 2 | 570 | 575 | 773 | 776 |
|  | 3 | 532 | 540 | 740 | 745 |
|  | **4** | **462** | **478\*** | **675** | **685\*** |
| **GLUT3** | 1 | 270 | 273 | 373 | 375 |
|  | 2 | 262 | 267 | 356 | 360 |
|  | 3 | 238 | 246 | 341 | 346 |
|  | **4** | **208** | **224\*** | **308** | **318\*** |
| **GLUT5** | 1 | 297 | 300 | 809 | 811 |
|  | 2 | 278 | 283 | 770 | 773 |
|  | 3 | 264 | 272 | 739 | 744 |
|  | **4** | **234** | **250\*** | **673** | **682\*** |

### 3.5 Mammalian and Avian GLUT Homology

**Table S6:** Comparison of known avian GLUT isoforms and their homology to humans. Data was aggregated from (M. S. Byers et al., 2017; Myrka & Welch, 2018; Sweazea & Braun, 2006; Kenneth C. Welch et al., 2013) and homology to humans was calculated using NCBI BLAST (Boratyn et al., 2012).

| GLUT | Localisation | Feature | Chicken to hummingbird sequence homology | Chicken to human sequence homology | Hummingbird to human sequence homology | Substrates (mammals) |
| --- | --- | --- | --- | --- | --- | --- |
| **GLUT1** | Ubiquitous | Basal glucose transport | 98% | 80% | 88% | Galactose, mannose, glucosamine |
| **GLUT2** | Liver, Pancreas, Intestine, Kidney | Insulin dependent | 89% | 65% | 64% | Fructose, Glucose, Galactose |
| **GLUT3** | Neurons, Liver, skeletal muscle | Insulin dependent | 87% | 70% | 73% | Glucose |
| **GLUT4** | Not found | Absence | N/A | N/A | N/A | Glucose |
| **GLUT5** | Intestine, brain, adipocytes, testes, skeletal muscle | Fructose transport | 81% | 64% | 66% | Fructose |

